# Assessment of emissions and potential occupational exposure to carbon monoxide during biowaste composting

**DOI:** 10.1101/2023.08.06.552181

**Authors:** Karolina Sobieraj, Karolina Giez, Jacek A. Koziel, Andrzej Białowiec

## Abstract

To date, only a few studies focused on the carbon monoxide (CO) production during waste composting; all targeted on CO inside piles. Here, the CO net emissions from compost piles and the assessment of worker’s occupational risk of exposure to CO at large-scale composting plants are shown for the first time. CO net emissions were measured at two plants processing green waste, sewage sludge, or undersize fraction of municipal solid waste. Effects of the location of piles (hermetised hall vs. open yard) and turning (before vs. after) were studied. Higher CO net emission rates were observed from piles located in a closed hall. The average CO flux before turning was 23.25 and 0.60 mg‧m^−2^‧h^−1^ for hermetised and open piles, respectively, while after – 69.38 and 5.11 mg‧m^−2^‧h^−1^. The maximum CO net emissions occurred after the compost was turned (1.7x to 13.7x higher than before turning). The top sections of hermetised piles had greater CO emissions compared to sides. Additionally, 5% of measurement points of hermetised piles switched to ‘CO sinks’. The 1-h concentration in hermetised composting hall can reach max. ∼50 mg CO·m^−3^ before turning, and >115 mg CO·m^−3^ after, exceeding the WHO thresholds for a 1-h and 15-min exposures, respectively.

## 1. Introduction

Concern for the environment has led to initiatives and changes in regulatory frameworks worldwide and especially in Europe. The need to manage growing amounts of organic waste (biowaste) resulted in a renewed interest in the aerobic biological processing. The availability of biodegradable waste and its particular presorted types continues to grow, and includes, inter alia, food and kitchen waste, garden waste, agricultural waste and sewage sludge [1]. Moreover, industrial waste (e.g., from papermaking processes) is also treated at full-scale composting plants.

The first large-scale European composting plants in the 1970s and 1980s, treated mainly unsorted municipal solid waste (MSW). Since then, major process improvements have been implemented [2]. In 2019, the European countries used composting as the predominant waste treatment method, and 60% of the total biowaste weight was treated in ∼3,400 facilities [3]. The new generation of composting plants has been managed with higher standards, including ‘best available technologies’ (BAT) [4]. One such standard requires hermitisation (i.e., enclosing compost piles indoors) to better control the process, improve the quality of the final product, and manage local emissions of odours and gaseous pollutants. However, hermitisation of composting raises concerns about the occupational health and safety for workers, due to emissions and accumulation of toxic gases, and inhalation exposure.

The composting process is a source of air pollutants, such as H_2_S, SO_2_, NH_3_, dust, odours, volatile organic compounds (VOCs), endotoxins produced by bacteria, protozoan parasites and allergic fungi [5]. Toxic air pollutants are generated during various compost process stages, and in addition to the management operations including storage, sorting, grinding and turning [6].

One of the least known toxic gases emitted from composting is carbon monoxide (CO). CO is classified as a major ambient air pollutant which has immediate negative effects on human health and life. Emerging body of research has shown CO presence during composting of the undersize fraction of municipal waste, agricultural waste, green waste or fruit and vegetables [7–12]. However, it is worth emphasizing, that the research conducted to date on CO production during composting concerned its distribution *within* the composted material [7,8,13]; the literature does not provide information on net CO emissions *from* the pile surface into air above.

To date, modelling of CO production during composting in a lab-scale closed reactor has shown that the CO concentration can reach 36.1% without ventilation and 3.2% when accumulated process gas is released daily [14]. If scaled up, such CO concentrations would greatly exceed the acceptable inhalation exposure limits established by the World Health Organization (WHO), set at a peak CO concentration of 90 ppm for 15 min of physical work [15]. In general, CO concentration of 100 ppm causes a headache, while further symptoms (e.g., nausea, dizziness, general malaise) emerge at 200-300 ppm [16]. Monitoring the CO exposure is, therefore, important as health effects can be misdiagnosed for other ailments, such as influenza or food poisoning [17]. The chronic CO inhalation at a lower concentrations can adversely affect the respiratory, circulatory and nervous systems [18].

To date, the extent to which composting plant workers are at risk due to CO inhalation is not known and more research is needed. Measurement of CO emissions from large compost piles is challenging due to inherent spatial and temporal variability. The static flux chamber method is the commonly used for measuring gas emissions from large surfaces. Originally derived from soil gas emissions studies, flux chamber method was adapted for anthropogenic emissions sources. The method is based on the use of static (non-flow-through) chambers [19]. For static chamber method, the increasing gas concentration as a function of time is used to back-calculate flux from the enclosed surface [20], as demonstrated for the flux of greenhouse gases such as N_2_O, CH_4_ or CO_2_ from soil [21]. In this research, the static flux chamber method was used for the operational simplicity needed for measurements at a large-scale plant.

Building on the research on CO production *inside* compost piles and aiming to bridge the knowledge gap in actual CO emissions *from* compost, we measured CO net emissions from surfaces of composted biowaste into air. To our knowledge, the CO net emissions assessment at large-scale composting plants was completed for the first time. This research was motivated by the need to assess the occupational risk of CO inhalation at composting plants and, if warranted, evaluate the need to implement the necessary safety measures. For this purpose, CO flux from compost piles was measured at two composting plants, one of which implemented current BAT guidelines for hermetisation. Effects of composting plant type (outdoors vs. enclosed indoors/hermetised) and compost pile turning were studied. Measured fluxes were used for modelling of potential occupational exposure to CO emissions.

## 2. Materials and Methods

### 2.1. Experiment matrix

Two composting plants representing differing technologies were selected. The first (Plant A) was in Rybnik, Poland, processing green waste (grass, leaves, branches) and sewage sludge from the “Boguszowice” wastewater treatment plant, 85 and 15% by fresh mass, respectively. Research focused on four compost piles located in the enclosed hall during September-October of 2021. The second (Plant B) was in Lubań, Poland, processing green waste (5 piles) and undersize fraction of municipal solid waste (1 pile), both in open yard.

Biowaste samples (approximately 10 kg each) were collected manually with a shovel from three random locations from every analysed pile. Each sample was then reduced to ∼0.7 kg using the quartering method. The age of the composting piles ranged from 1 to 4 weeks (Plant A) and 4 to 8 weeks (Plant B). Experimental matrix is summarized in Table 1.

**Table 1.**
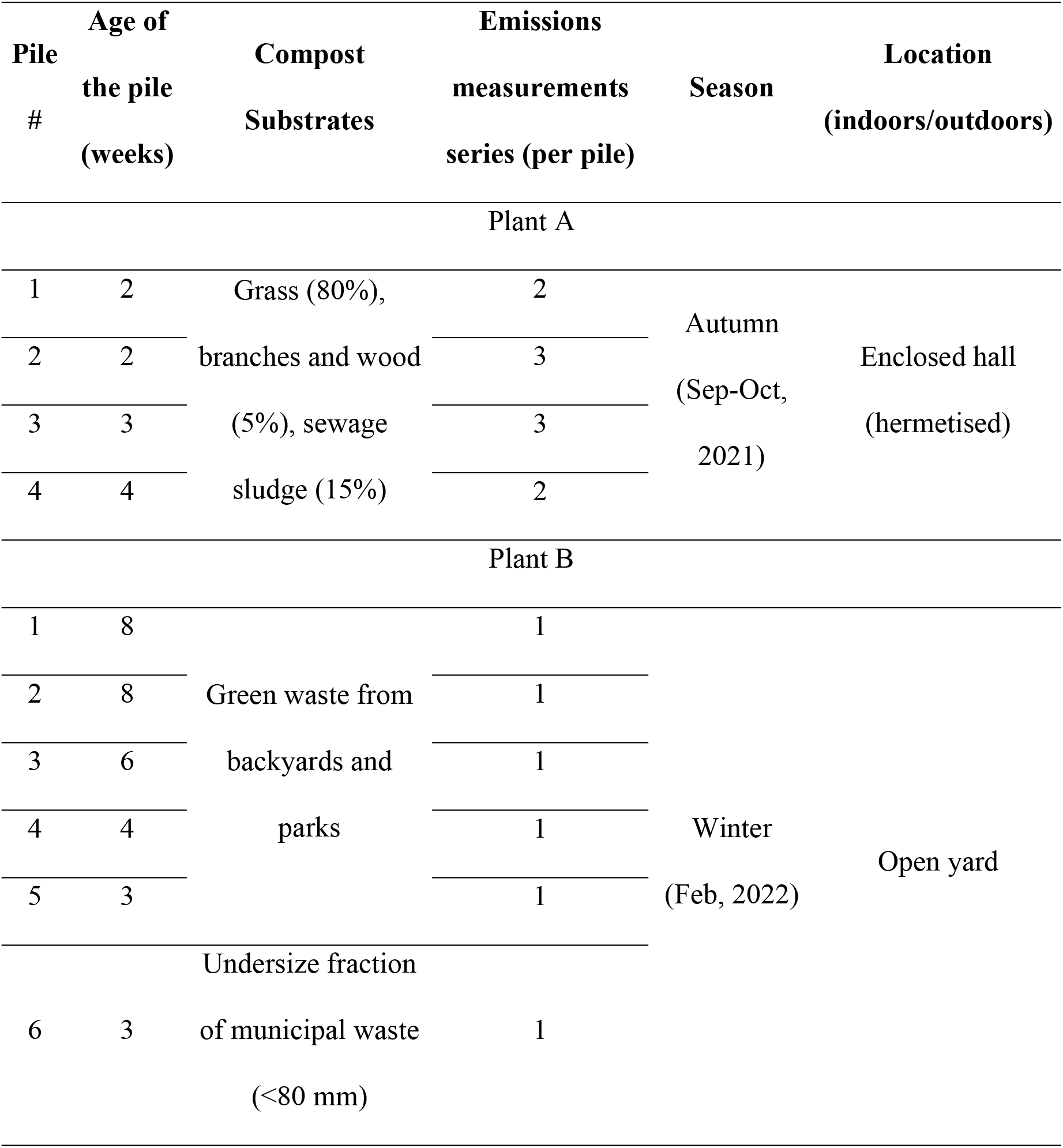
Experiment matrix for CO emissions measurements from biowaste compost.

### 2.2. Biowaste characterization

Samples were analysed for the dry matter content in accordance with PN-EN 14346:2011 [22], at 105 °C with RadWag WPT/R C2 (Radom, Poland) with an accuracy of 0.01 g and thermal testing chamber KBC-65 (WAMED, Warsaw, Poland). The organic dry matter content was determined according to PN-EN 15169:2011 [23] at 550 °C using the muffle furnace Snol 8.1/1100 (Utena, Lithuania). The respiratory activity (AT_4_) was measured as an indicator for compost stability using OxiTop Control system (WTW, Weilheim in Oberbayern, Germany) in accordance with [24].

### 2.3. Analysis of process gas emissions from compost piles

The measurement of CO emissions from compost piles was performed using the flux chamber method [25]. A plastic box with a volume of 0.071 m^3^ was adapted to serve as a flux chamber. Two valves were installed onto the chamber, one for gas sampling and the other for pressure equalization. Gas sampling valve enabled connection with the Kimo KIGAZ 300 gas analyser (Sauermann-KIMO Instruments, France) via a silicone tube (Fig 1) and CO concentration measurement (ppm). Ancillary measurements of CO_2_ and O_2_ was also conducted as they are considered co-dependent with CO [8,9]. Internal chamber temperature was measured (± 0.1°C) with a thermocouple.

**Figure 1.**
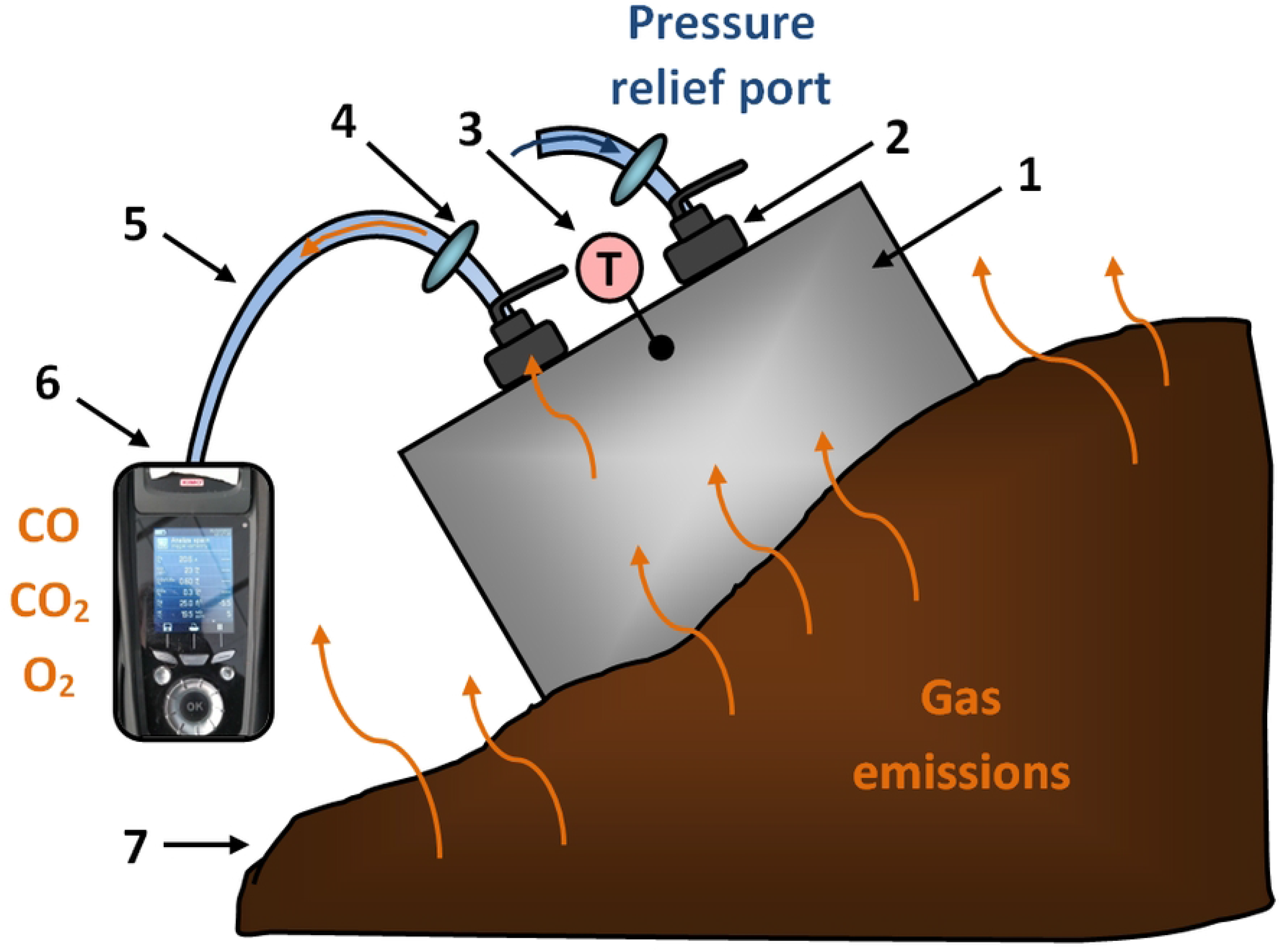
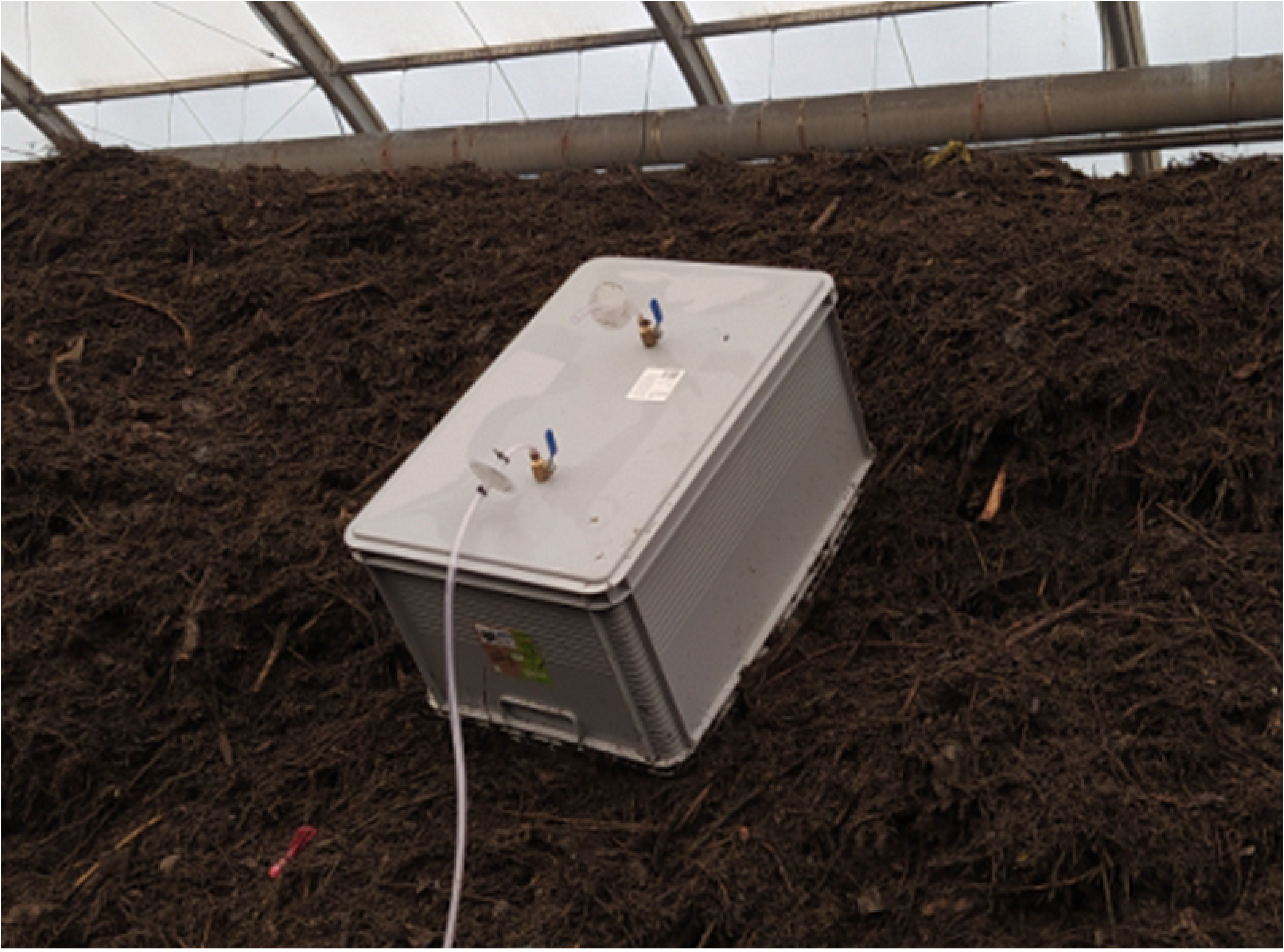
Flux chamber sampling of CO emissions. a) cross-sectional schematic, 1 – flux chamber, 2 – valves, 3 – thermocouple, 4 – purification filter, 5 – silicon tube, 6 – gas analyser, 7 – composting pile; b) flux chamber enclosing emitting surface of a green waste pile in hermetised composting Plant A

The flux chamber was placed on each pile in three locations along its length, on both sides and in its top (total of n = 9; D1-D9) according to the scheme (Fig 2). Due to the difficult access to pile 6 in plant B, measurements were made only for D1-D3. To improve the enclosure of emitting surface during the measurement, the chamber was pounded into the pile or, in the case of a more homogenous material, gently pressed into the pile. The gas analyser was equipped with an internal pump (1 L‧min^−1^) which facilitated real-time concentrations measurement. After connecting and calibrating of the gas analyser, and placement on the pile surface, each measurement was carried out for 5 min, and its course (changes in CO concentration over time, ppm) was recorded with a camera (Xiaomi Redmi Note 8T, Beijing, China). Data was then processed manually, by entering real-time concentrations every 5 s into spreadsheet (summarized in Supplementary Material). After each measurement, the analyser was disconnected from the chamber and there was a short pause to flush remaining sampled gas and return to the ambient atmospheric levels (CO ∼0 ppm, O_2_ ∼20.2%, CO_2_ ∼0%). Each measurement series (Table 1) was done once a day and included measurements of CO, CO_2_ and O_2_ concentrations *before* and *after* pile turning. Daily turning during initial phase of composting was facilitated by a self-propelled turner for windrows.

**Figure 2.**
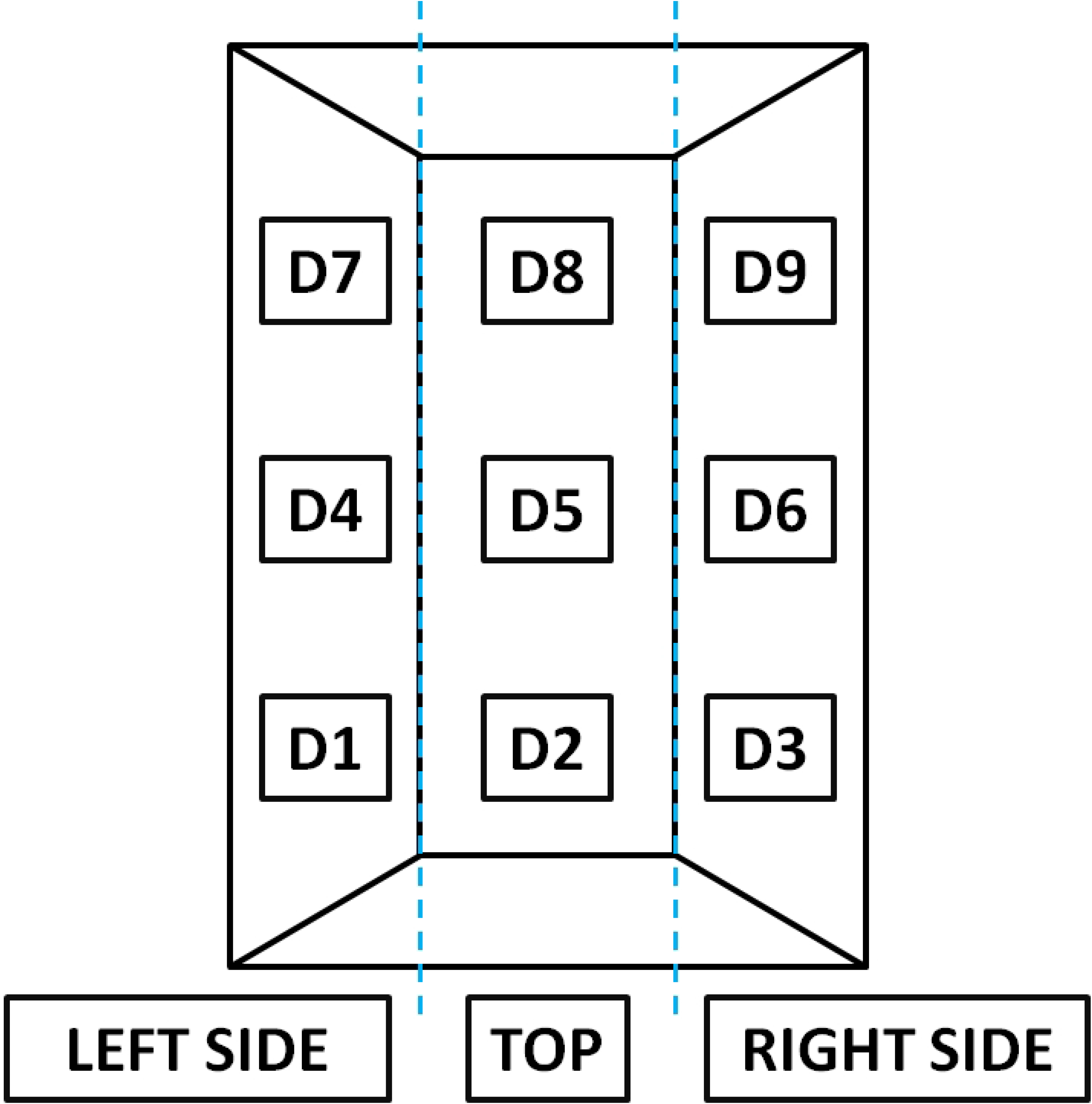
The top view of composting pile with the location of flux chamber placement for CO emissions measurements. Locations D1, D4, and D7 represent left side of the pile; D2, D5, and D8 – pile tops, while D3, D6, and D9 represent pile right side

### 2.4. Estimating CO emissions

The UK Environmental Agency’s methodology (LFTGN07 Guidance on monitoring landfill gas surface emissions) [25] was adopted for estimating CO emissions. Measured CO concentrations (ppmv) were converted to mass/volume units (mg‧m^−3^) at standard temperature and pressure (273 K and 101.3 kPa) using:

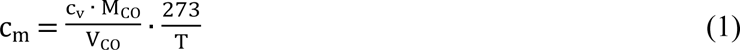

where:

c_m_ – CO concentration, mg‧m^−3^,

c_v_ – CO concentration, ppmv,

M_CO_ – molecular weight of CO, M_CO_ = 28×10^3^ mg‧mol^−1^,

V_CO_ – molecular volume of CO at standard conditions, V_CO_ = 0.0224 m^3^‧mol^−1^,

T – analysed gas temperature during measurement.

CO flux for each measurement location (D1-D9 on compost pile, Fig 2) was calculated using:

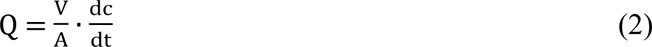

where:

Q – CO flux, mg‧m^−2^‧s^−1^,

V – volume of the flux chamber, V = 0.071 m^3^,

A – emitting surface area of compost pile enclosed by the flux chamber (flux chamber footprint), A = 0.23 m^2^,

dc/dt – rate of change of measured CO concentration in the flux chamber with time, determined by plotting CO concentrations on chart with the x-axis representing time (s) and the y-axis representing the mass concentrations (mg‧m^−3^), mg‧m^−3^‧s^−1^.

### 2.5. Modelling of CO emissions in the composting plant

The modelling of CO emissions during 1 h of operation of the enclosed (hermetised; airtight) composting hall with a 1,000 m^3^ of headspace, with a total area of piles of ∼1200 m^2^ was performed. The 1 h period was chosen for modelling due to the average worker time for turning one pile, and therefore 60 min of exposure to CO emissions per pile. The ‘worst-case-scenario’ was assumed, i.e., no ventilation in the composting hall and CO emissions allowed to accumulate. The mass of emitted CO during t = 1 h for both ‘before’ and ‘after’ turning of the compost material was:

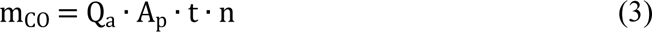

where:

m_CO_ – mass of the emitted CO during t = 1 h for both before and after compost turning, mg,

Q_a_ – averaged flux of CO from measurement locations D1-D9 on compost pile, mg‧m^−2^‧s^−1^,

A_p_ – surface of n=1 compost pile, A_p_ = 300 m^2^,

t – time, t = 3600 s,

n – number of piles inside hermetised composting hall, n = 4.

CO concentration in the headspace of the composting hall after t = 1 h accumulation in both ‘before’ and ‘after’ compost turning scenarios was:

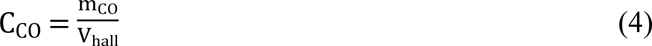

where:

C_CO_ – CO concentration in the headspace of the composting hall after accumulation for t = 1 h for both before & after compost turning, mg‧m^−3^,

V_hall_ – volume of the headspace of the airtight composting hall, V_hall_ = 1,000 m^3^.

CO concentration in the headspace of the composting hall after accumulation (C_CO_) was then converted to the ppm values:

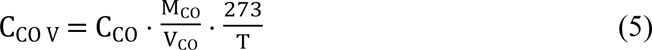

where:

C_CO v_ – CO concentration in the headspace of the composting hall after accumulation for t = 1 h for both before/after compost turning, ppm.

### 2.6. Statistical Analyses

All data were analysed using Statistica StatSoft Inc., TIBCO Software Inc, i.e., estimating the measurements mean, standard deviation, conducting the correlation analyses between CO emissions and CO_2_, O_2_ concentration and temperature.

## 3. Results

### 3.1. Compost biowaste characterization

Compost piles in Plant A (hermetised) were characterized by similar dry matter content (DM) and dry organic matter (OM) content (DMO) (Fig 3). The DM values were ∼35% and ranged from 34.9% (pile 1) to 36.6% in pile 4. For DMO, the highest mean value was noted for pile 1 (66.8% DM), and the lowest (61.4% DM), was obtained for pile 2. Different biowaste properties were observed at Plant B (open yard), where DM varied from 69.8% in case of pile 6 to 34.6% for pile 1. DMO levels ranged from 26.6% DM to over 50% DM. Clearly, the process parameters were more difficult to control in an open yard operation.

**Figure 3.**
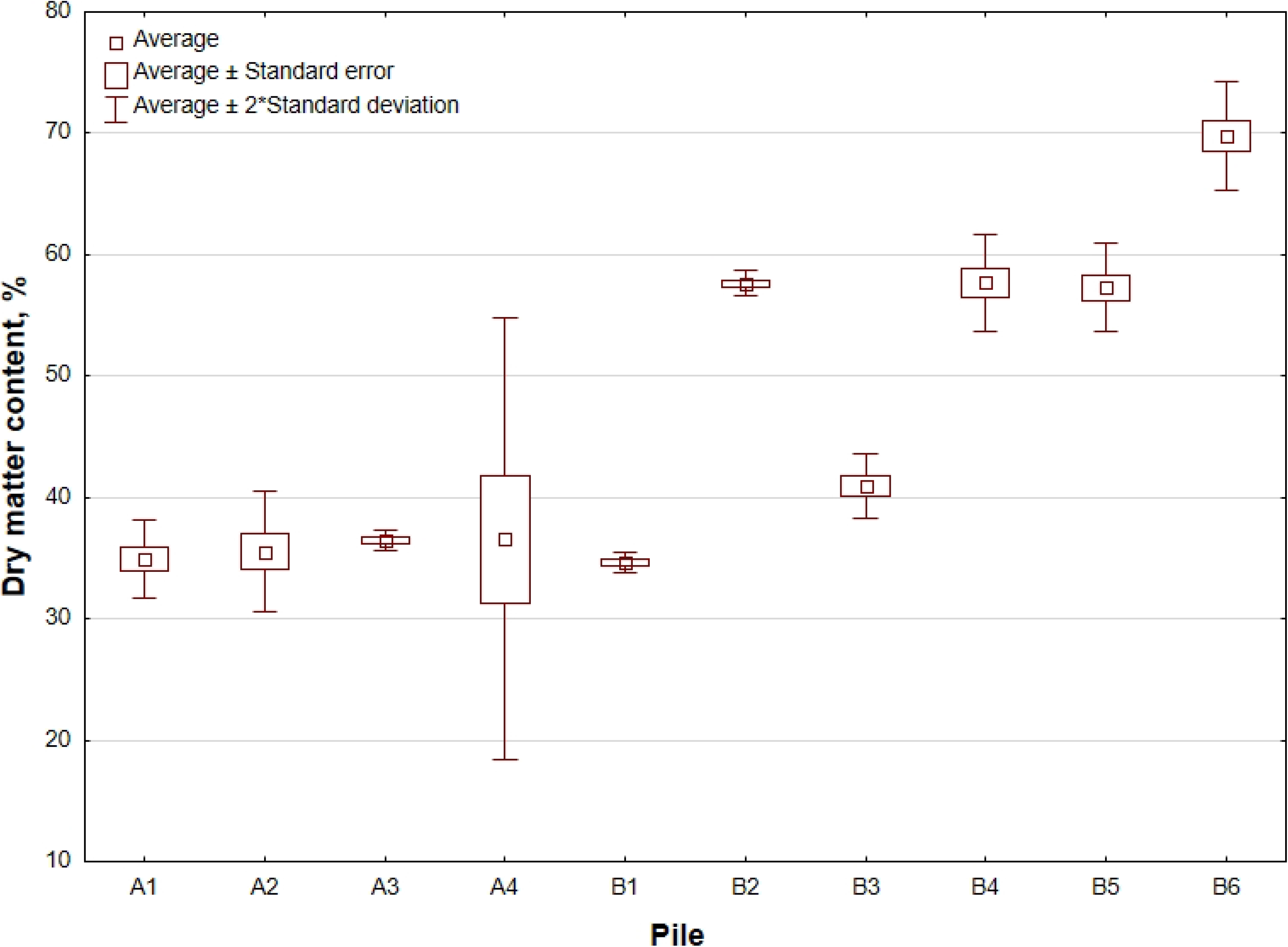
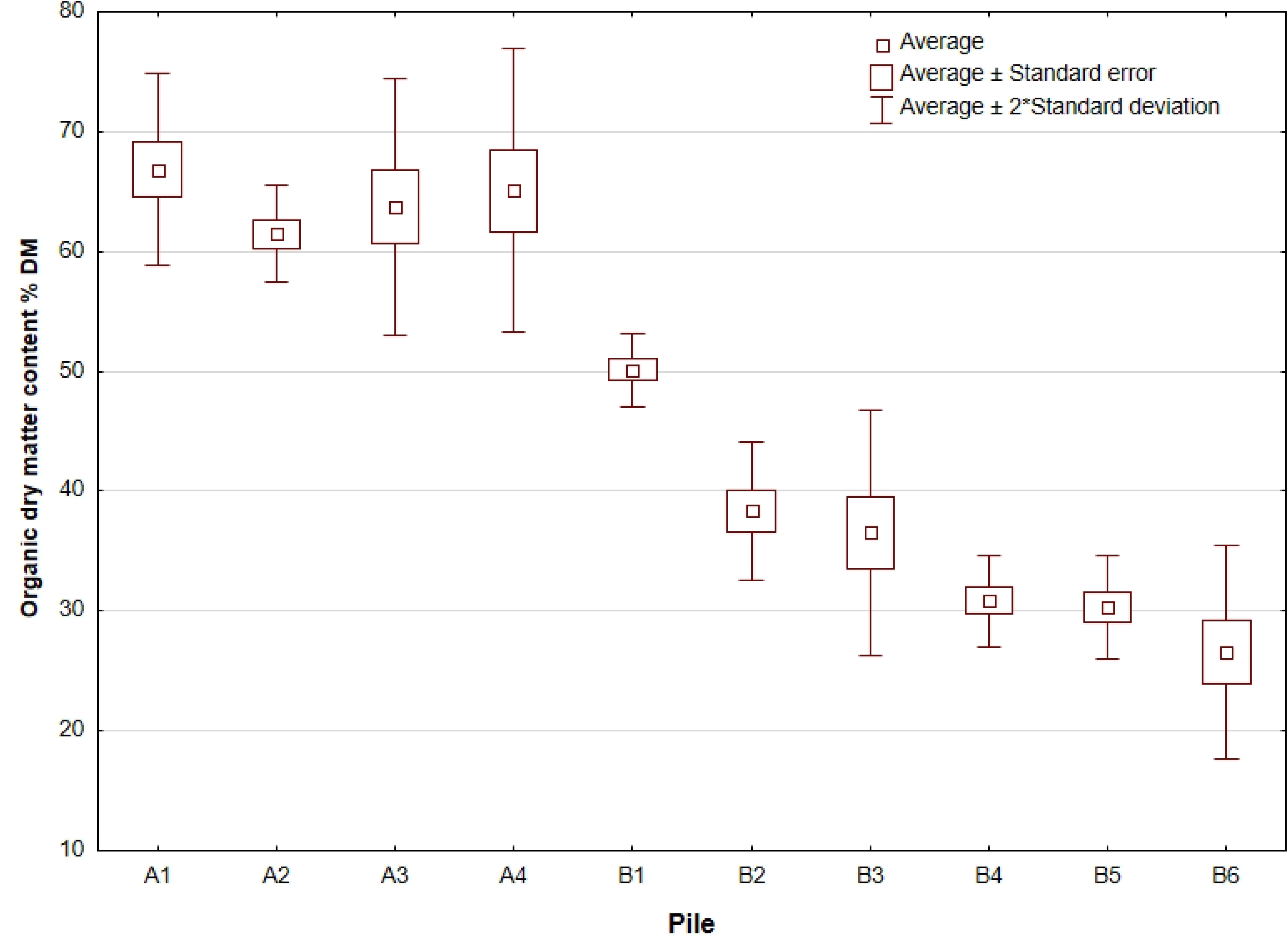
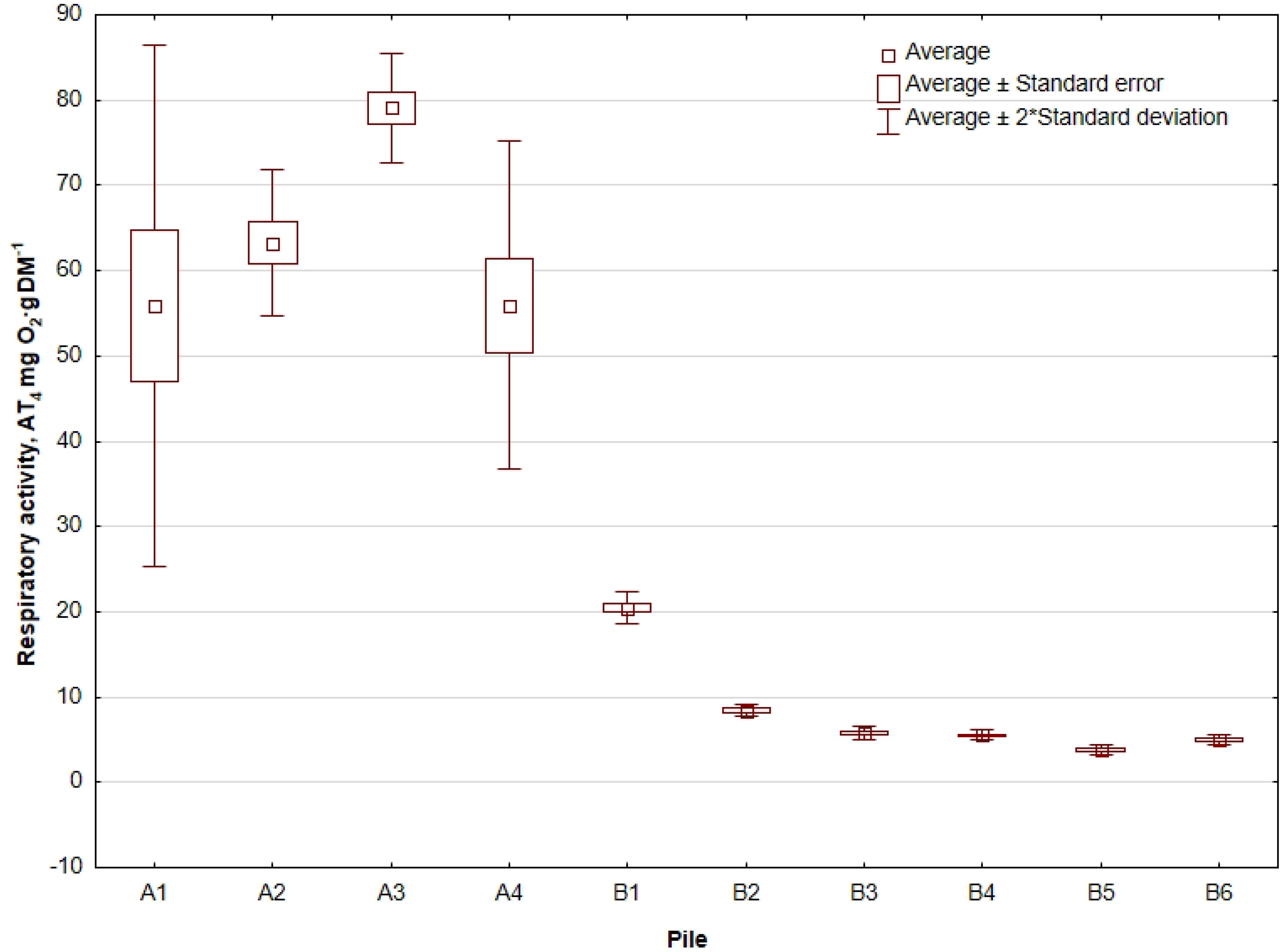
Compost properties for piles 1-4 in Plant A (hermitised, A1-A4) and Plant B (open yard, B1-B6) a) dry matter content, %; b) organic dry matter content, % D.M.; c) respiratory activity AT_4_, mg O_2_·g DM^−1^

The respiratory activity was different for piles in Plants A and B. In general, the compost in Plant B (open yard) can be classified as stabilized material (AT_4_ <10 mg O_2_‧g DM^−1^) [26]. The exception was pile 1, for which the AT_4_ > 20 mg O_2_‧g DM^−1^. In turn, Plant A (hermetised) piles were characterized by high respiratory activity where the limit value for stabilized compost was exceeded, and AT_4_ ranged from 52.3 to as high as 80.3 mg O_2_‧g DM^−1^.

### 3.2. CO fluxes from composting piles

The assessment of CO net emissions at large-scale composting plants was completed. Detailed measured CO concentrations and CO flux estimations are summarized in Excel spreadsheets in Supplementary Materials. Tables 2 and 3 summarize the spatial distribution of CO flux from piles before and after turning, in a hermetised and open yard plants, respectively.

**Table 2.**
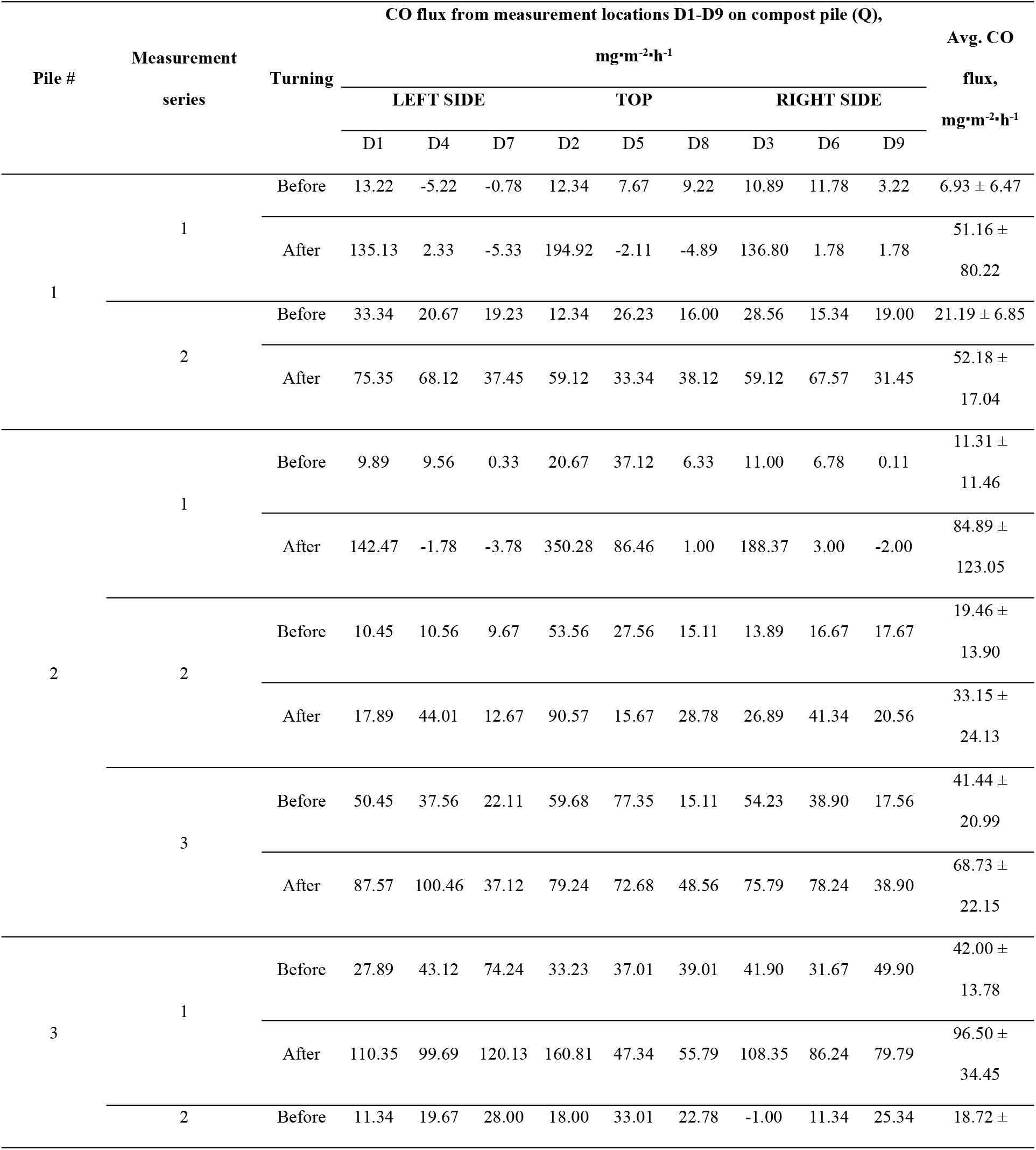
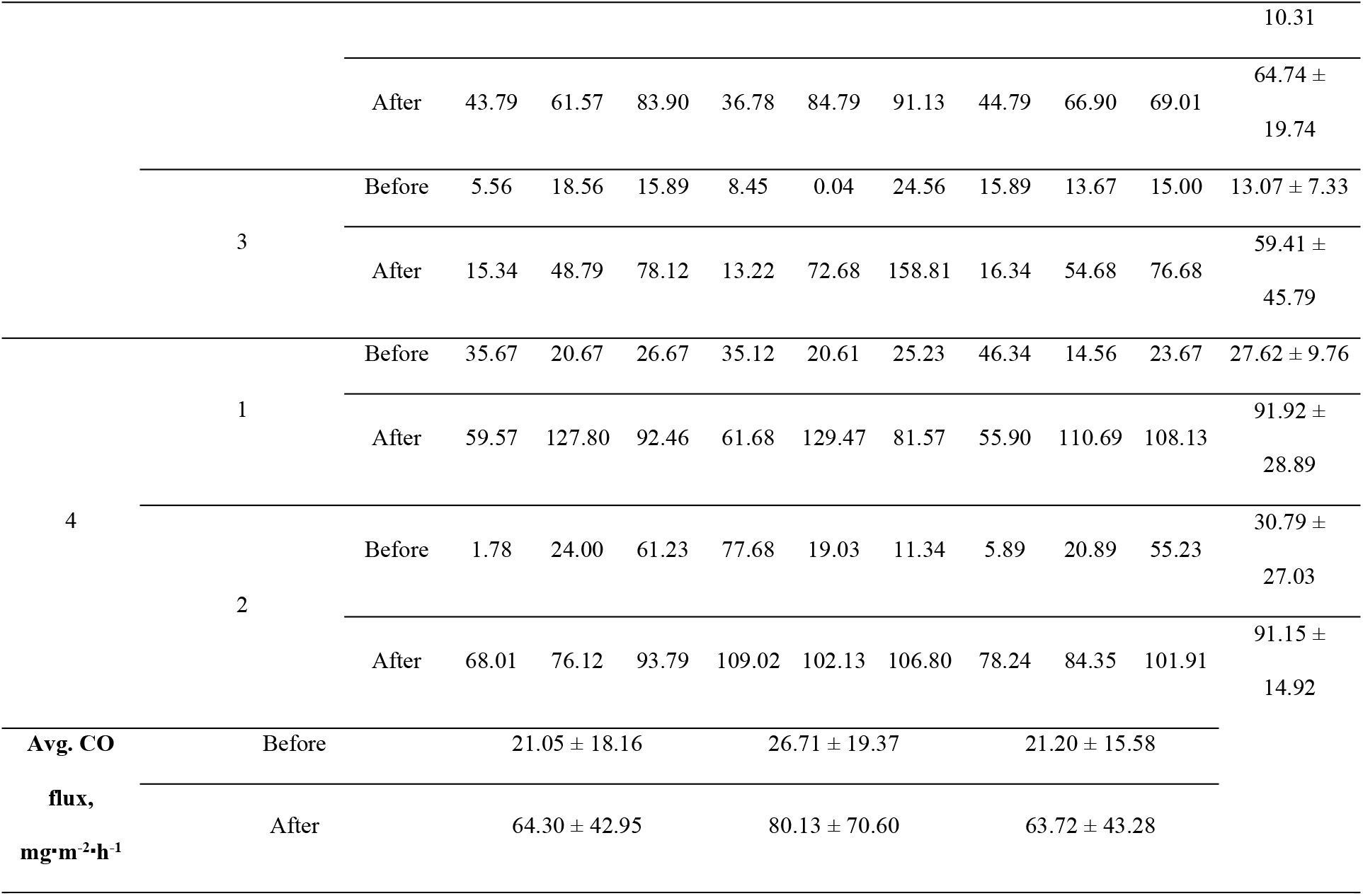
Spatial distribution of CO flux (Q) from compost piles in hermetised plant (Plant A) before and after turning.

**Table 3.**
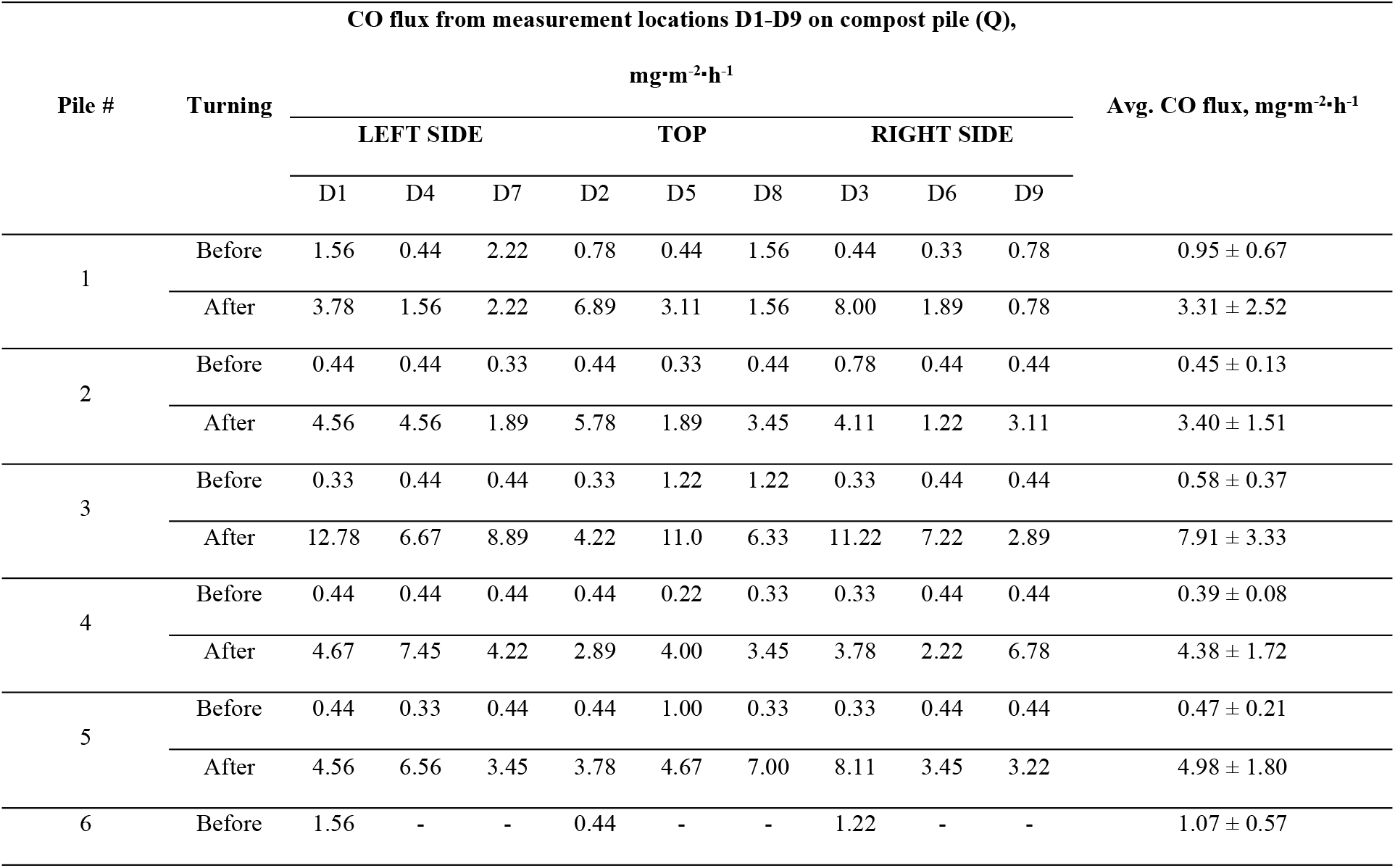
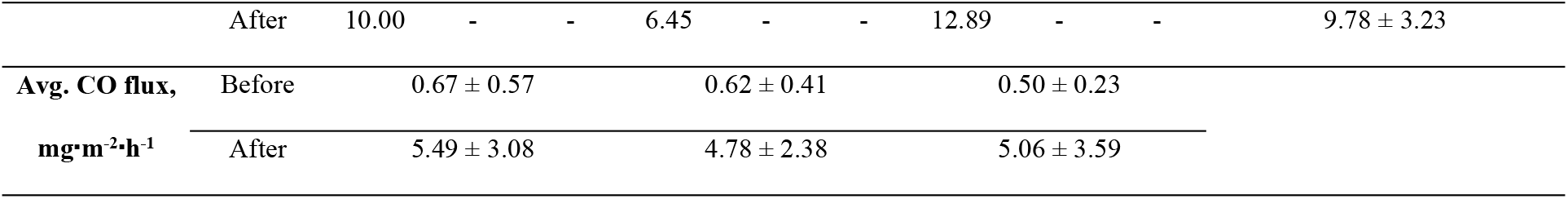
Spatial distribution of CO flux (Q) from compost piles in open-yard plant (Plant B) before and after turning.

At Plant A, a higher average CO flux was measured at the top of the piles, compared with CO flux from the sides. The CO flux was 26.71 mg CO‧m^−2^‧h^−1^ vs. 21.05 and 21.20 mg CO‧m^−2^‧h^−1^, and 80.13 mg CO‧m^−2^‧h^−1^ vs. 64.30 and 63.72 mg CO‧m^−2^‧h^−1^ for top, left and right side of the piles before and after turning, respectively (Table 2). At Plant B, the left side of the pile was emitting more CO compared with the top (Table 3). The highest CO fluxes here were measured on the left side of the piles (0.67 and 5.49 mg CO‧m^−2^‧h^−1^, before and after turning respectively).

Piles in hermetised hall generated more CO emissions than those outdoors, both before and after compost turning. The average CO flux in all cases was higher after the material was turned; the increase varied from 1.7x to 7.4x for plant A (hermetised, Tab. 2) and from 3.5x to 13.7x for plant B (open yard, Table 3). The lowest recorded average CO flux was 6.93 and 0.39 mg CO‧m^−2^‧h^−1^, while the highest reached ∼100 and ∼10 mg CO‧m^−2^‧h^−1^ (with max. values equal to 350 and 12. 9 mg CO‧m^−2^‧h^−1^, values for plant A and B, respectively).

Importantly, a negative CO flux was recorded at 9 measurement points in hermetised plant A (5% of total measurement locations, Table 2). In most cases, negative CO fluxes were observed after material turning (points D7, D5, and D8 for pile 1, measurement series 1; D4, D7, and D9 for pile 2, series 2). The CO sinks were not distributed evenly, i.e., most of them were located at the sides of the piles (>50% ‘CO sinks’ occurred on the left side and two of them on the right). The strongest ‘CO sink’ achieved −5.22 mg CO‧m^−2^‧h^−1^ (point D4 in pile 1 before turning, series 1), while the weakest – −0.78 mg CO‧m^−2^‧h^−1^ (point D7, the same pile).

Based on the data presented in Tables 2 and 3, overall net CO emission factors for hermetised and open composting piles were developed (Table 4) for the before and after turning for both hermetised and open plants. The average CO flux was lower before the compost is turned. In the ‘before turning’ scenario it reached 23.25 and 0.60 mg CO‧m^−2^‧h^−1^ for Plants A and B, respectively, and 69.4 and 5.11 mg CO‧m^−2^‧h^−1^ after the turning. The before/after turning ratio was higher for hermetised piles (0.34 vs. 0.12 for piles located outdoors). However, the range of before/after ratios was broad. For hermetised plant it ranged from negative (−5.37) up to >6, while for open yard piles it ranged from 0.03 to 1.00.

**Table 4.**
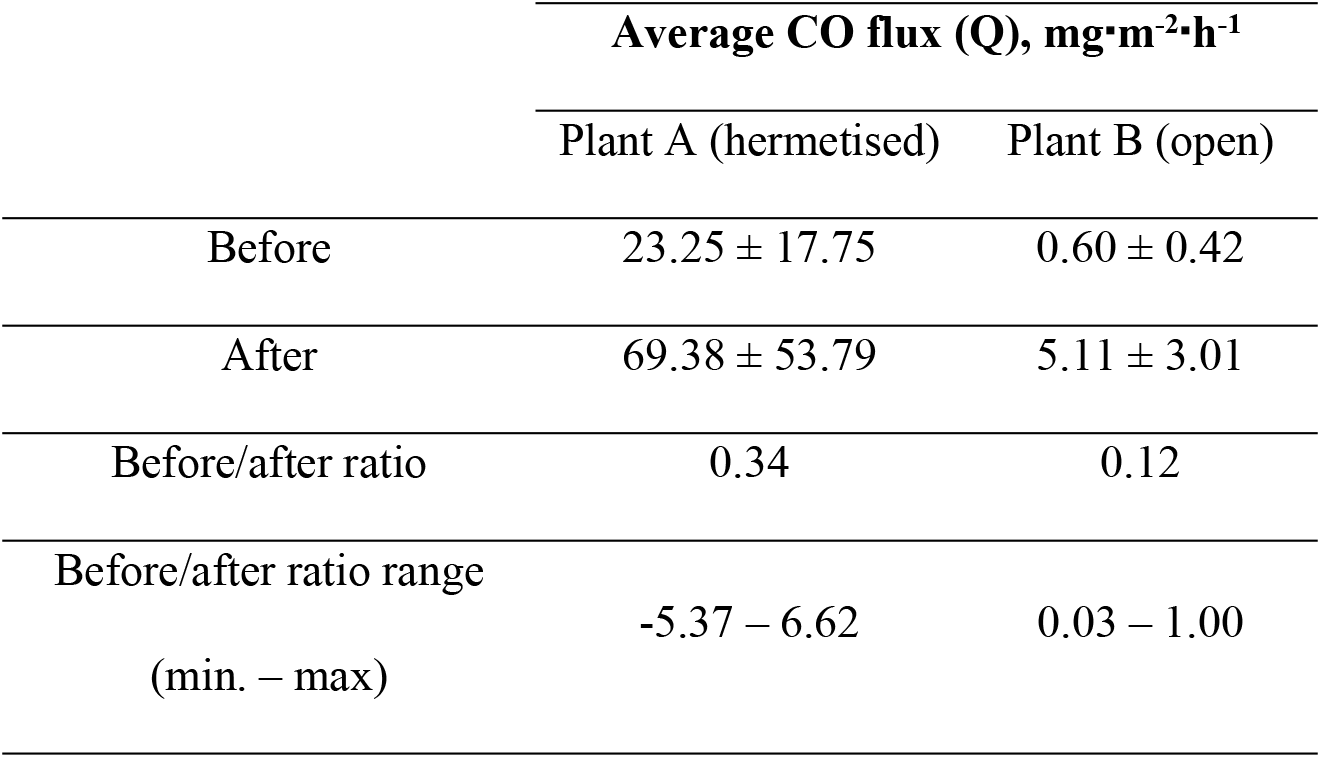
Summary of averaged CO fluxes for hermetised (Plant A) and open (Plant B) piles Average CO flux (Q), mg‧m^−2^‧h^−1^.

### 3.3. CO concentration accumulation in the hermetised composting plant

The modelling of CO emissions during 1 h of operation of the enclosed (hermetised) composting hall with a cubature of headspace 1,000 m^3^, processing green waste with an annual capacity of 60,000 Mg (one-time area of piles in the hall ∼1,200 m^2^) was performed. Modelling has shown that the concentration of accumulated CO in the hall headspace during 1 h in ‘before turning’ scenario can reach from 8.3 to even 50.4 mg‧m^−3^ (Table 5). In each of the analyzed cases, this concentration increased after turning the material, reaching values from 1.7x to over 7x higher, i.e., raising concerns about the potential occupational risk during a typical 1 h-long pile turning. In the ‘after turning’ scenario, CO levels in the hall headspace after 1 h reached >60 mg‧m^−3^, exceeding 100 mg‧m^−3^ in 4 analyzed cases. The maximum modelled CO concentration was 110.3 mg‧m^−3^.

**Table 5.**
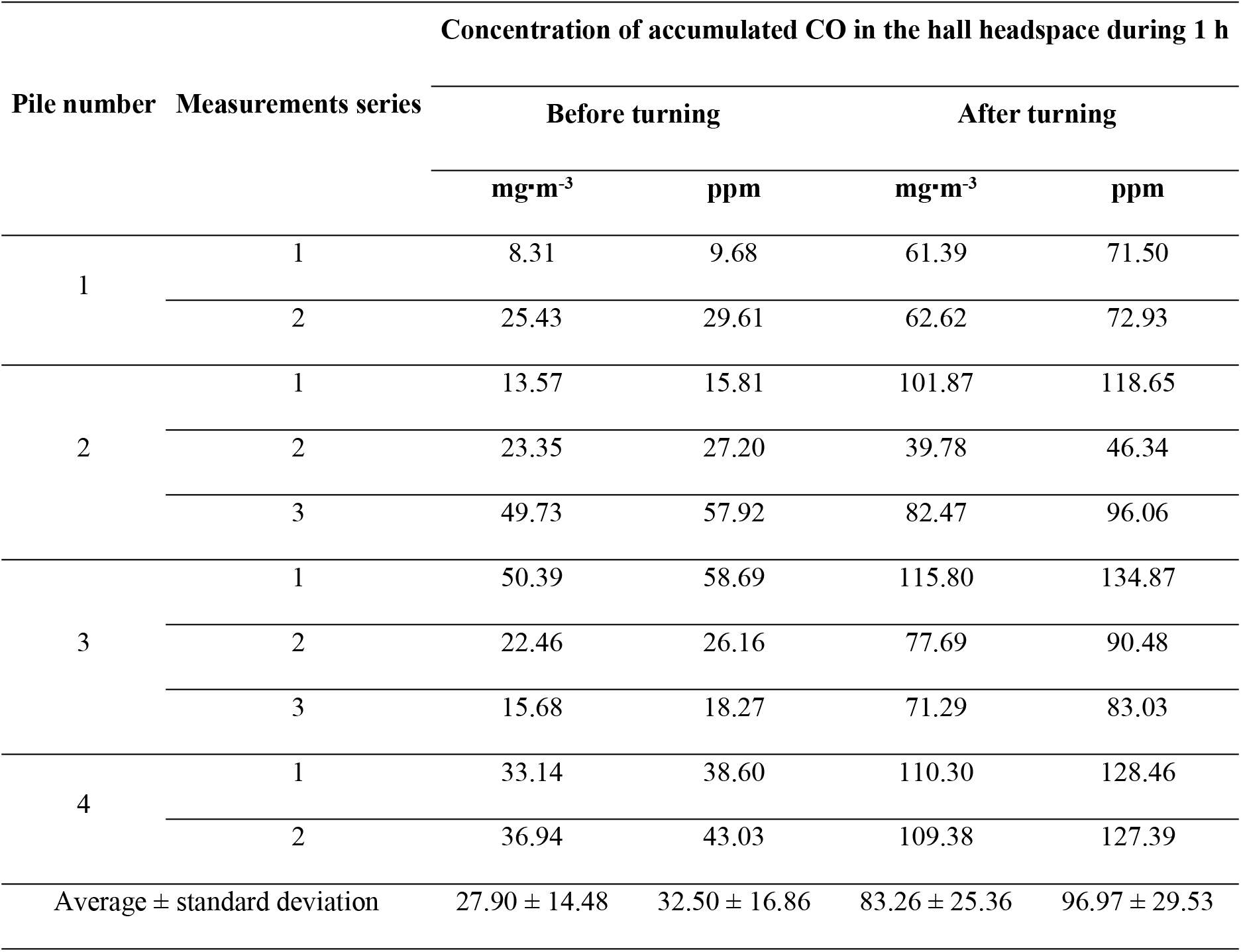
Concentration of accumulated CO in the hall headspace during 1 h modelled for hermetised plant.

### 3.4. Relationship between CO and other process gases and temperature

Correlation analysis showed that in Plant A (hermetised) CO emissions followed measured CO_2_ concentrations (Pearson correlation coefficient r ranged from 0.55 to 0.91) and negative correlation with measured O_2_ concentration (r ranging from −0.78 to −0.91, Table 6), both before and after turning. This is in contrast to the observations of other researchers, reporting that the increased availability of O_2_ stimulates the production of CO related to thermal degradation of OM [7,9]. No statistically significant correlations between those gases were obtained for Plant B (open yard, Table 7). More research is needed to evaluate the kinetics of CO, CO_2_ and O_2_ as the effect of turning and its frequency.

**Table 6.**
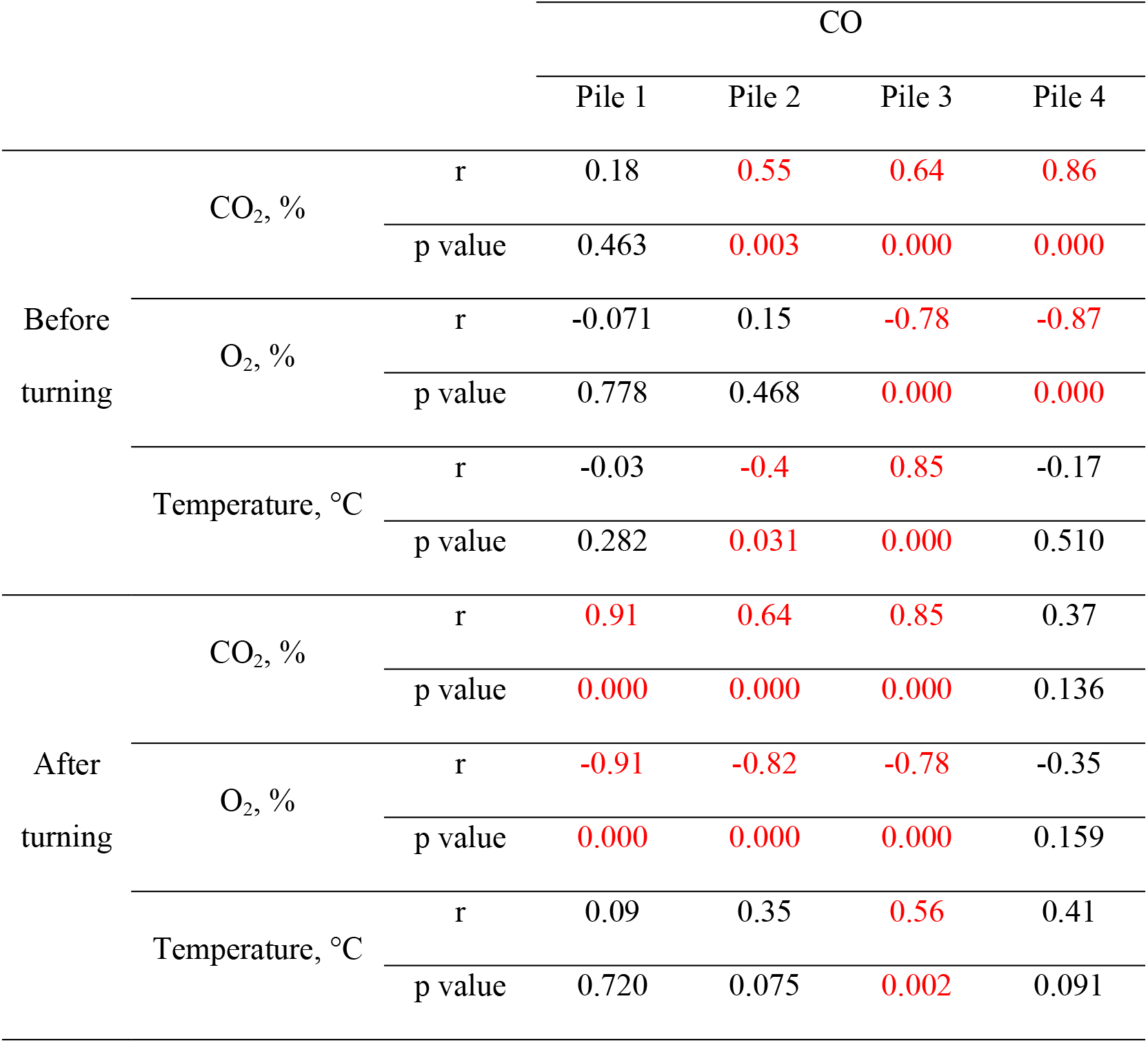
Correlation between CO and other process gases and temperature in Plant A (hermetised) for a probability level of α=0,05. statistically significant correlation coefficients are marked in red, r – Pearson correlation coefficient

**Table 7.**
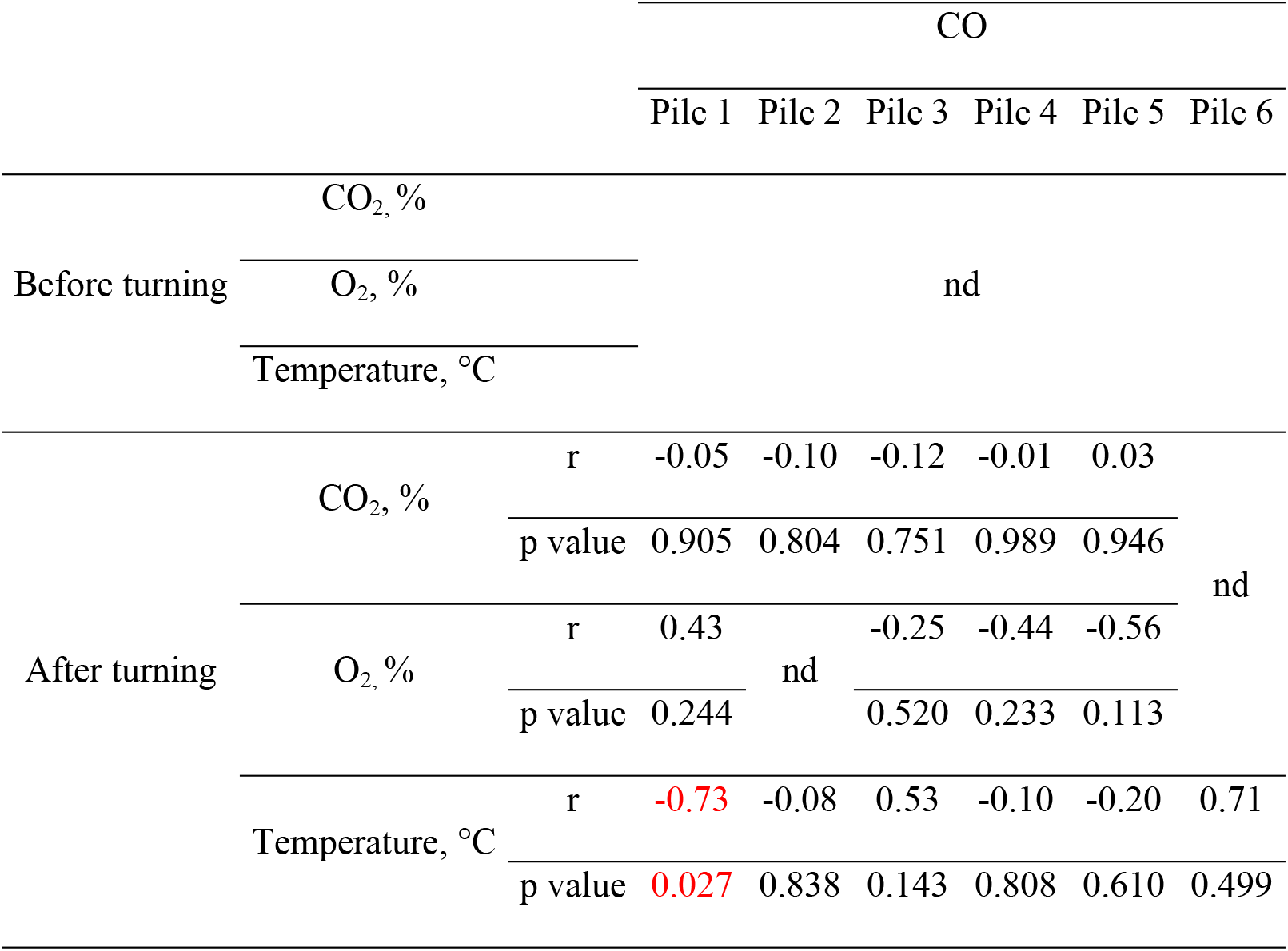
Correlation between CO and other process gases and temperature in Plant B (open) for a probability level of α=0,05. statistically significant correlation coefficients are marked in red, r – Pearson correlation coefficient, nd – no data

There was no statistically significant correlation between CO emissions and temperature, as observed by other researchers [9,10,13,27]; the only exceptions were pile 2 (before turning) and pile 3 (before and after turning) in Plant A and pile 1 in plant B (after turning). However, the data obtained was inconsistent; for pile 2 and pile 1 (Plant A and B, respectively) the correlation was negative, in the first case the r was low (−0.42, Table 6), while in the case of pile 3 (plant A) the correlation was strongly positive (r equal to 0.85 and 0.56 before and after turning the material, respectively).

## 4. Discussion

To date, only a few studies focused on the CO production during waste composting; all were targeted on CO inside piles. Here, data of CO net emission from compost piles is shown for the first time. The comparison of process-based CO emissions for ‘before’ and ‘after’ compost turning is important both in terms of occupational safety, and for improved inventory of CO sources in local and regional air quality.

Regarding the occupational safety, the topic of CO emissions accumulation in enclosed spaces is rarely discussed in the context of waste management. Related studies were conducted mainly in relation to the storage of wheat, rape, wood pellets or during the processing of such materials, e.g. wood drying, in rooms similar in nature to closed composting halls [28,29]. For the first two, emission factors reached up to 200 mg CO‧ton^−1^ (rape) and 9 mg CO‧ton^−1^ (wheat grain) per day. Moreover, the recorded CO levels in the storage and processing of wood materials exceeded the permissible values for warehouses [30].

According to the study conducted here, the CO accumulation in hermetised compost halls should also be of concern. Based on emissions modelling, averaged CO level before turning reached nearly 30 mg CO‧m^−3^, and after – more than 80 mg CO‧m^−3^, with single values exceeding 50 and 100 mg CO‧m^−3^, respectively. According to WHO guidelines, the 30 mg CO‧m^−3^ should not be exceeded during 1-h work and 100 mg CO‧m^−3^ during 15 min of moderate physical activity [31]. This is important because of the toxic CO impact on human health. Prolonged exposure to CO causes the formation of carbohydrate hemoglobin (COHb) due to the higher affinity of CO for hemoglobin compared to O_2_ [31].

The *duration* of the high CO concentrations in hermitised plans is also important in the context of the exposure of composting plant workers. Composting facilities often work continuously with three 8-h shifts. A typical worker repeats scheduled turning of piles over entire shift, and thus, may be exposed to increased CO emissions throughout the entire 8 h of work. The initial phase of exposure to CO starts with the first pile turning. COHb concentration increases rapidly at the beginning of exposure to a constant CO concentration [31]. Stabilization takes place after 3 h, and the steady state, when the CO concentration in alveolar breath and ambient air is ∼equal, is achieved after 6-8 h, i.e., practically during one work shift in a closed composting hall [32]. Moreover, high CO levels may be present in closed halls for a longer period, even several months during cool season when the ventilation is low. During the research on emissions from wood pellets, the CO concentration was equal to 21 mg‧m^−3^ even after 3 months from the beginning of storage of this raw material [30]. This is particularly important due to the fact that long-term exposure to lower CO levels results in much greater health impact than short-term exposure to high concentrations of this gas. The health consequences of chronic CO exposure include, inter alia, heart failure, asthma, stroke, tuberculosis, pneumonia, cognitive memory deficits or sensorimotor changes [15]. Human activity level during exposure to CO is also important. Considering that compost plan workers of the composting plants sometimes handle waste manually, it should be taken into account that in combination with long shifts in hermetised environment with high levels of CO and potentially other highly toxic gases such as H_2_S, and moderate-to-high activity (and therefore inhalation rate) pose synergistically elevated risks.

Moreover, the CO levels may increase again during composting with increasing ambient temperature [30]. The peaks of higher CO concentration were observed after 100 days from the start of the process, when the temperature reached 80 °C [27]. This means that in the context of exposure of workers to the negative effects of CO, monitoring should be carried out throughout the process, not only in its initial stage. In addition, it is possible that piles originally considered as ‘safe’ (with lower CO net emissions), such as those processed outdoor in Plant B, when moved to a composting hall with more favourable thermal conditions, may again exhibit higher CO emissions.

Taking into account the spatial variability of gaseous emissions from compost piles, the CO gradient distribution indicates that its level is higher in top of the piles [27]. This is confirmed by the observations made for hermetised Plant A, where ∼1.2x higher CO fluxes, both before and after turning, were measured at the top of piles. A similar situation was also noted during the storage of wood pellets [30]; the highest values, significantly exceeding the permissible levels of CO emissions, were recorded at the top of the pile. It was also noted in case of other pollutants emission, such as VOCs and N_2_O [27,33]. This tendency is related to the so-called ‘chimney effect’ in the pile, which is caused by the temperature profile within the material and occurs as a result of convection [33,34]. In this way, the warmer gas migrates from the core of the pile due to buoyancy leaves it through the top, while the cooler air enters the sides of the pile, close to the ground [35]. The chimney effect was observed in this research for CO emissons from the pile. This is important from occupational safety of plant employees who work with pile levelling. Additionally, CO, being slightly lighter than air, rises in the enclosed hall and accumulates in its upper part [15]. Thus, high-off-the-ground cabin location of common machinery (excavators, turners, or shredders) may result in greater risk to operators exposure to CO emitted from the top of the piles. On the other hand, the chimney effect was not noted in the case of open piles in Plant B, where the higher average CO flux occurred on the left side of the pile. This may be related to the influence of external conditions, such as wind direction. This is confirmed by research conducted by [27], who explain the asymmetric distribution of process gases in the pile with higher pressure and pore gas dilution in the area of the pile not sheltered from the wind.

It should be emphasized that compost can not only be a ‘source’ but also a ‘sink’ of CO, which in hermetised plant occurred in 5% of flux measurement locations. Emerging evidence have shown that CO production during composting has a twofold character and is based on (1) the activity of microorganisms (biotic CO production), and on (2) thermochemical processes dependent on temperature and O_2_ concentration (abiotic CO production) [9]. Furthermore, when the CO production is biotic, net CO emission is the result of the CO formation by bacteria and its metabolism (microbial oxidation); the enzyme carbon monoxide dehydrogenase (CODH) plays a key role controlling both processes [36]. The same situation was observed with soils [36]; early research dating back to the 1970s identified soils not only as a CO producers, but also as the main sinks of atmospheric CO [37]. The nature of CO uptake is mainly based on microbial activity, as confirmed by studies of autoclaved soil and the use of antibiotics [37–39]. For this reason, CO consumption is also limited by the concentration – an increased level of CO can inhibit the metabolism of bacteria. An important element of the biotic CO uptake studied for soils is also the fact that these processes occurred under both aerobic and anaerobic conditions [38]. This issue becomes important in the context of studies on aerobic and anaerobic bacteria functioning in an environment with >1% CO concentration, which use the enzyme carbon monoxide dehydrogenase (CODH) to metabolize CO [40]. Due to the bidirectional activity of this enzyme, enabling the reversible process of CO oxidation to CO_2_, it can be hypothesized that, apart from bacteria that only produce/consume CO, there are also strains that carry out both of these processes. The responsibility of microorganisms for ‘CO sinks’ in composting piles in this research may also affect the spatial distribution of spots with negative CO fluxes. About 78% of them occurred on both sides of the piles, creating chimney effect of CO uptake on the pile sides and emission of CO from the top of the pile. Since CO and O_2_ concentration were positive correlated, this effect could be caused by the transfer of aerobic CO-metabolizing microorganisms from sites with less nutrient availability to areas with higher O_2_ concentration and decomposable OM content.

The second aspect of this study, i.e., the determination of CO net emission factors from open and hermetised piles before and after turning (Tab. 5) is needed for atmospheric air quality modelling and CO source inventories. Open yard Plant B had a much lower CO emission potential compared with hermetised Plant A. However, according to the research conducted by [30], the outdoor composted material emits most of the gases in warm season. The authors associated this with the close correlation of CO concentration and temperature, which is especially visible in thermophilic conditions [30]. During present study, CO fluxes from open piles were estimated in winter, when the ambient temperatures were low. It is also worth noting that no statistically significant correlation between CO concentration and temperature was observed. However, it should be remembered that the dependence of CO production on temperature refers to the thermal conditions *inside* the composted material [8]. The temperature measured in these studies prevailed in flux chamber headspace, i.e., directly above the pile. Considering the ambient conditions (low temperatures in winter), it can be assumed that the temperature in the flux chamber correspond to the conditions under which CO was *net emitted*.

## 5. Conclusions and recommendations

Research on CO net emissions from biowaste composting on industrial scale has shown its dependence on turning and plant type (open yard vs. hermetised). Higher CO net emission rates were observed for piles located in an enclosed composting hall, separated from ambient conditions (23.25 and 69.38 mg CO‧m^−2^‧h^−1^ before and after turning, respectively). In each of the analyzed cases, maximum CO emissions occurred after compost turning. The areas with increased CO emissions for hermetised piles were the tops with ‘CO sinks’ spots on the sides, showing the ‘chimney effect’ of CO distribution. Modelling of CO emissions during 1-h of work in a closed hall has shown that it can reach max. ∼50 mg CO·m^−3^ (59 ppm) before turning, and >115 mg CO·m^−3^ (135 ppm) after, exceeding the WHO thresholds for an 1-h and 15-min exposures, respectively.

The results show that due to the nature of work in composting plants (operating machine with cabins high above ground, occasional manual labour, 8-h shifts), personal protective equipment should be implemented for workers exposed to CO emissions (e.g., personal CO detectors, appropriate breathing masks with filters). This is especially important for people working with biowaste turning or manual levelling on top of piles. Additionally, it is recommended that the time spent in the closed composting hall be shortened to a minimum and limiting activities to moderate physical effort. Access to composting halls should be limited only to authorized persons, equipped with appropriate safety equipment, and following protocols. Automating turning and eliminating workers exposure could be developed and implemented to the composting practice. Due to the CO tendency to accumulate in the upper part of halls, it is also recommended to install alarms, especially above compost piles. Since CO emissions are variable and may increase with the temperature, reaching several peaks throughout the process, it is recommended to monitor it continuously throughout the composting process, not only in its initial stage. Engineering design should consider adequate ventilation for operations involving human operators.

Since this study has shown that compost can be considered not only as a ‘producer’, but also as a ‘sink’ of CO, based on studies on CO consumption conducted for soils, it can be hypothesized that during bio-waste composting aerobic and anaerobic bacteria are responsible for the CO uptake, possibly using the CODH enzyme to metabolize CO. Further research identifying the mechanisms of biotic CO uptake should be conducted as a future strategy for CO emission mitigation.

## 6. Author Contributions

**Karolina Sobieraj**: Conceptualization, Data curation, Formal analysis, Funding acquisition, Methodology, Visualization, Writing – original draft; **Karolina Giez**: Investigation, Funding acquisition, Project administration, Resources; **Jacek A. Koziel**: Data curation, Formal analysis, Supervision, Writing – review & editing. **Andrzej Białowiec**: Conceptualization, Supervision, Validation, Writing – review & editing.

## 7. Acknowledgements

The presented article results were obtained as part of the activity of the leading research team – Waste and Biomass Valorization Group (WBVG).

## 8. Funding

The research was financed under the individual student research project “Młode umysły – Young Minds Project” (project title: “Study of CO emissions from turned piles on a technical scale and exposure of composting plant workers to CO emissions”, number: N010/0007/21) from the subsidy increased for the period 2020–2026 in the amount of 2% of the subsidy referred to Art. 387 (3) of the Law of 20 July 2018 on Higher Education and Science, obtained in 2019.

The publication was financed by the project “UPWR 2.0: international and interdisciplinary programme of development of Wrocław University of Environmental and Life Sciences”, co-financed by the European Social Fund under the Operational Program Knowledge Education Development.

## 9. Supporting information

All measured CO concentrations, CO fluxes estimation according to the UK Environmental Agency’s methodology and modelling of CO concentration in closed composting hall can be found in the Supplementary Material.

## Notes

### Competing Interest Statement

The authors have declared no competing interest.

